# Low aflatoxin levels and flowering delay in *Aspergillus flavus*-resistant maize lines are correlated with increased corn earworm damage and enhanced seed fumonisin content

**DOI:** 10.1101/2020.02.03.933309

**Authors:** Subbaiah Chalivendra, Fangneng Huang, Mark Busman, W. Paul Williams, Jong Hyun Ham, Geromy G. Moore

## Abstract

Preharvest mycotoxin contamination of field-grown crops is influenced not only by the host genotype, but also inoculum load, insect pressure and their confounding interactions with seasonal weather. In two field trials, we observed a preferred natural infestation of specific maize (*Zea mays* L.) genotypes by corn earworm (*Helicoverpa zea* Boddie) and investigated this unexpected interaction. These studies involved four maize lines with contrasting levels of resistance to *Aspergillus flavus*. The resistant lines had 7 to 14-fold greater infested ears than the susceptible lines. However, seed aflatoxin B_1_ levels, in mock- or *A. flavus*-inoculated ears were consistent with maize genotype resistance to *A. flavus*. Further, the corn earworm-infested ears had greater levels of fumonisin content in seeds than uninfested ears, indicating that the insect may have vectored native *Fusarium verticillioides* inoculum. The two maize lines with heavy infestation showed delayed flowering. The availability of young silk for egg-laying could have been a factor in the pervasive corn earworm damage of these lines. At the same time, *H. zea* larvae reared on AF-infused diet showed decreasing mass with increasing AF and >30% lethality at 250 ppb. In contrast, corn earworm was tolerant to fumonisin with no significant loss in mass even at 100 ppm, implicating the low seed aflatoxin content as a predominant factor for the prevalence of corn earworm infestation and the associated fumonisin contamination in *A. flavus* resistant lines. These results highlight the need for integrated strategies targeting mycotoxigenic fungi and their insect vectors to enhance the safety of crop commodities.

**IMPORTANCE:** *Aspergillus* and *Fusarium* spp. not only cause ear rots in maize leading to crop loss, they can also contaminate the grain with carcinogenic mycotoxins. Incorporation of genetic resistance into breeding lines is an ideal solution for mycotoxin mitigation. However, the goal is fraught by a major problem. Resistance for AF or FUM accumulation is quantitative and contributed by several loci with small effects. Our work reveals that host phenology (flowering time) and insect vector-mycotoxin interactions can further confound breeding efforts. A host genotype even with demonstrable resistance can become vulnerable due to seasonal variation in flowering time or an outbreak of chewing insects. Incorporation of resistance to a single mycotoxin accumulation and not pairing it with insect resistance may not adequately ensure food safety. Diverse strategies including host-induced silencing of genes essential for fungal and insect pest colonization and broad-spectrum biocontrol systems need to be considered for robust mycotoxin mitigation.

## INTRODUCTION

Besides causing crop damage and economic loss to the grower, mycotoxigenic fungi pose a serious risk to human and livestock health due to the contamination of commodities with carcinogenic and neurotoxic secondary metabolites known as mycotoxins. Aflatoxin B_1_ (**AF**) is the most dangerous among mycotoxins due to its very potent carcinogenicity. *Aspergillus flavus*, an opportunistic pathogen, is the predominant species that contaminates cereal and oil seed crops with AF. Although not as genotoxic as AF, fumonisins (**FUM**) are associated with esophageal cancer, particularly due to cytotoxicity of fumonisin B_1_ (**FB_1_**). They are also among the most common food- and feed-contaminating mycotoxins in many countries (BIOMIN Mycotoxin Survey 2015). FUM are produced by *Fusarium* species*, F. verticillioides* (formerly known as *F. moniliforme*) being the predominant contaminant of commodities (Munkvold 2003). *A. flavus* and *F. verticillioides* cause ear rots in maize (*Zea mays* L.), a globally important food, feed and fuel crop of high productivity. Co-contamination of commodities with AF and FUM has been reported, particularly, in high cancer-risk areas (Sun et al. 2011; Shirima et al. 2013; Guo et al. 2017). Studies in animal models indicate an additive or even synergistic effect on liver cancer due to an exposure to both mycotoxins (Lopez-Garcia 1998; World Health Organization 2018).

Aspergillus and Fusarium ear rots are more frequent in warmer and drier cropping seasons or a warmer and wetter weather combination at the time of harvest, and are often exacerbated by insect damage. Insect-vectored inoculum can breach the natural plant defense. The invasive methods of inoculation by chewing and piercing insects would bypass resistance mechanisms, such as remote defense signals triggered in the husk, silk or seed surface in response to natural infection. Consequently, ear rot diseases are more common in the southern United States (US) and lowland tropics (Miller, 1994; reviewed in Cotty and Jaime-Garcia 2007; Santiago et al. 2015). Among insect pests infesting maize, European Corn Borer (**ECB**) causes the most serious damage (Boyd and Bailey, 2001; Hutchison et al. 2010). It not only injures plants, exposing them to infection, but also vectors ear rot and stalk rot fungi, particularly *F. verticillioides* and *F. graminearum* (Widstrom 1992). Extensive use of **Bt** (*Bacillus thurigiensis Cry*stal proteins-expressing) maize with its high efficacy against ECB, has reduced overall ECB populations in the US (Hutchison et al. 2010). Maize pests previously considered as secondary to ECB are now taking its position (Bowers et al. 2014). Corn earworm [**CEW**; *Helicoverpa zea* (Boddie); formerly in the genus *Heliothis*] has become the most economically important pest in the southern United States where non-freezing winters are conducive for CEW to multiply by 4-7 generations in a year. Resistance of this pest to a wide range of insecticides and to Bt maize has also been documented (Capinera 2004; Dively et al. 2016; Kaur et al. 2019). Although CEW has multiple crop and weed hosts, maize is its preferred host (Johnson et al. 1975). Annual yield loss due to CEW ranges from 2-17% for field corn and up to 50% in sweetcorn in the southern US. *A. flavus* and *F. verticillioides* invade the seed through silk and are also vectored by CEW and other ear-infesting insects (Munkvold and White 2016). *F. verticillioides* can grow also as an endophyte through root or stem infection, and is vectored also by insects such as ECB that feed on vegetative tissues (Blacutt et al. 2018). Unlike a strong association observed in the case of FUM contamination (e.g., Smeltzer 1959; Dowd 2000; Mesterházy et al. 2012), seed AF levels were poorly correlated with CEW damage caused by either natural invasion (Ni et al. 2011) or manual infestation (Lillehoj et al. 1984). A meta-analysis of published work showed a 59% reduction in the mean FB_1_ concentration in Bt maize compared to the non-Bt control (Cappelle 2018). A complete mitigation of AF or FUM, requires control of multiple pests, including CEW (Abbas et al. 2013; Bowers et al. 2014; Porter and Bynum 2018).

In addition to facilitating fungal colonization, insect infestation can also enhance mycotoxin production in host tissues (Döll et al. 2013; Drakulic et al. 2015, 2016). In turn, mycotoxigenic fungi can affect insect vector infestation by inducing volatile production in host tissues. This is particularly well documented in the case of *Fusarium* species (Schulthess et al. 2002; Piesik et al. 2011; Drakulic et al. 2016). For example, pre-inoculation of maize with *F. verticillioides* was shown to enhance the fecundity and rate of development in Lepidopteran and Coleopteran pests (Ako et al. 2003), while retarding larval development in western corn rootworm (*Diabrotica virgifera virgifera*; Kurtz et al. 2010). We observed a preferential CEW infestation and increased FUM contamination in *A. flavus* resistant maize lines in our field trials. This previously unreported or overlooked observation was pursued to unravel the factors underlying this novel host-pathogen-insect interaction. Although late flowering might have facilitated enhanced oviposition by *H. zea* in these maize lines, our analysis suggests that the toxicity of AF to CEW is a more compelling reason for the observed prevalence of ear damage in the low AF-accumulating genotypes.

## RESULTS

### Unusual weather pattern and corn earworm outbreak in 2018 summer

During the summer of 2018, daily profiles of rain fall and air temperature patterns were different from past years’ average in Louisiana as well as many of the maize-growing states in US. The growing season was shorter (late April to early August) due to extended cold temperatures into the beginning of the planting season and relatively warmer and drier days during the early crop growth period (**Fig. S1**). April 2018 was the coldest April month since 1997 based on US average temperatures (and for Iowa and Wisconsin, it was the coldest April since records began in 1895). In contrast, May 2018 was the hottest May on record, breaking the record set in May 1934 during the Dust Bowl (National Oceanic and Atmospheric Administration: https://www.noaa.gov/). The unseasonal and steep warming after protracted cold seems to have favored an explosion of CEW population as indicated by a heavy infestation of ears in both of our experimental plots. CEW incidence was also reported from maize fields in other states in southern (Porter and Bynum 2018) as well as northern US (e.g., Handley 2018). In spite of two applications of a strong broad-spectrum insecticide before and after silking, the insecticide seems to have failed to reach silks covered by the husks. Further, all ears were bagged immediately after inoculation/pollination, which concealed earworm damage until developing ears were sampled for analysis.

### CEW infestation was significantly greater in *A. flavus* resistant maize lines

During sampling of ears later in the season (July), we noticed that the two resistant lines, the hybrid Mp313ExMp717 and the inbred CML322 showed greater infestation by CEW than the susceptible lines GA209xT173 and B73 (**Fig. 1**, left panels). The infestation was <10% in susceptible lines and it ranged from 22% to 68% in the resistant lines. The maize lines used in the two field trials have been extensively validated in the field and are often used as checks for evaluating new genotypes and in mapping resistance loci (e.g., Mideros et al. 2012; Guo et al. 2017). Despite our concerns that the distinctive patterns of CEW infestation might potentially interfere with the genetic response of maize lines to *A. flavus*, AF measurements showed that the genotype responses were robust in spite of CEW infestation. As described in the **MATERIALS AND METHODS** section, we harvested and utilized all ears in the plots to obtain robust AF data. The insect infestation was 8-fold greater in CML322 than observed in B73 ears in the mock-inoculated set. Inoculation with the highly toxigenic Tox4 strain resulted in a significant (p<0.01) and nearly 4-fold decrease in the infestation of CML322, but still 2-fold greater than infestation in B73. This is inversely correlated with >3-fold increase in seed AF content in Tox4-inoculated CML322 ears. As expected from its susceptibility to *A. flavus* colonization, B73 seeds accumulated >100 ppb of AF even in mock-inoculated (Control) ears and >500 ppb in Tox4-inoculated ears. These AF levels are >12-19 fold higher than those measured in CML322 seeds (**Fig. 1B**, right panel). CEW infestation was also greater in the resistant hybrid (Mp313E x Mp717) than in the susceptible hybrid by >30-fold in the control set and by 7-fold in the inoculated set (**Fig. 1A**, left panel). Infestation was inversely correlated with seed AF levels in hybrids as well. The susceptible hybrid (GA209×T173) had 100 ppb in control seeds and >400 ppb of AF in the inoculated set (i.e., 3 and 24-fold greater than in the resistant hybrid). Unlike the resistant inbred CML322, the resistant hybrid showed no difference in either AF content or CEW infestation between the control and CA14-inoculated ears. Analysis of variance (ANOVA) confirmed that only the host genotype (i.e., resistance to *A. flavus*) affected infestation highly significantly (>99.99% confidence level) and inoculation-induced differences were not statistically different (**Table S1 and S2**).

**Fig. 1.**
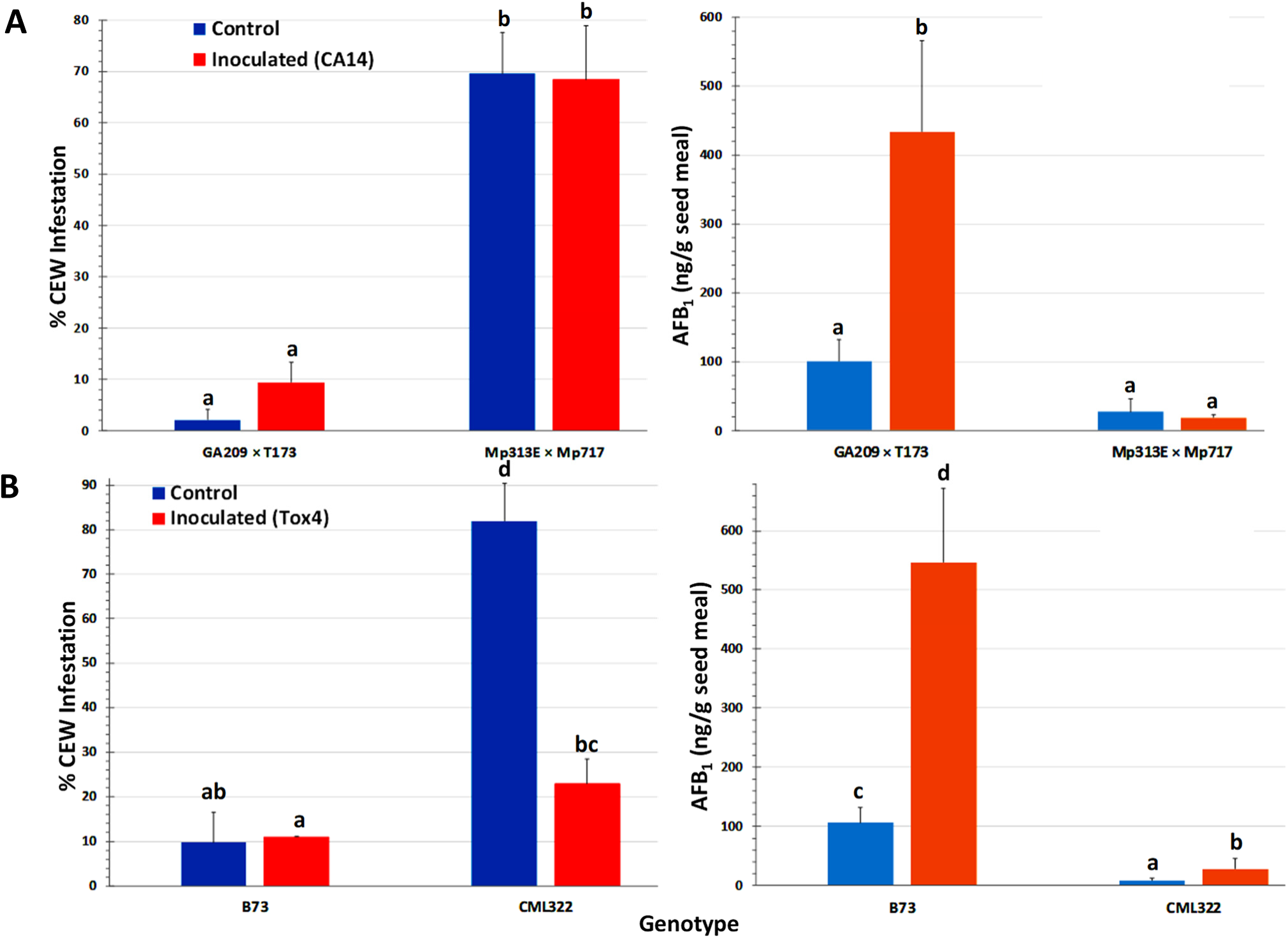
Rate of corn earworm infestation (left panels) and seed AF content (right panels) in maize lines. (**A**) Data is from hybrid plots. Infestation was significantly dependent on the host genotype with very little difference between control (mock-inoculated) and CA14-inoculated set. Seed AF content in CA14-inoculated set and the control were also similar in the resistant hybrid (Mp313E x Mp717). (**B**) Data shown is from inbreds. There was a similar negative relationship between CEW infestation rate and seed AF content as was observed in hybrids. Infestation was significantly dependent on the host genotype with very little difference between control (mock-inoculated) and Tox4-inoculated plots except in the case of CML322. The resistant inbred showed only 30% infestation in Tox4 inoculated set compared to the control. Seed AF levels were significantly higher in B73 both in control and inoculated ears than those of CML322. Values shown are average + SE. Significant differences (P value <0.05) between each data set were tested using an ANOVA (Supplemental Table 1) followed by Tukey’s multiple-comparisons post hoc test (Supplemental Table 2) in R (version 3.6.2). Means are significantly different if marked by a different letter.

### CEW infestation is negatively correlated with seed AF content

Not surprisingly, ANOVA of AF content revealed that the host genotype and inoculation with toxigenic *A. flavus* strains showed highly significant independent (or direct) as well as interaction effects on seed AF content. As indicated by the data presented in Fig. 1, infestation was also significantly related to AF content, although the interaction effect of genotype with infestation on AF was not significant (**Tables S3 and S4**). Both the resistant genotypes (CML322 and Mp313E×Mp717) manifested robust resistance to *A. flavus* and accumulated less than 30 ppb of AF in the seed either in the control (via colonization of native *A. flavus* strains) or the inoculated set. Conversely, the susceptible inbred and hybrid accumulated 100 and 500 ppb in control and inoculated sets, respectively. AF content is inversely correlated with CEW infestation pattern in each of the four maize genotypes. This relationship becomes clear when the data is combined for control and inoculated sets in each genotype (**Fig. 2)** or when all data is combined (**Fig. S2**). It is of interest to note that the uninfected controls from both resistant lines showed a numerical but statistically insignificant increase in AF in CEW-infested ears. AF was scarcely detectable levels in the uninfested and uninoculated controls (a mean value of 6 ppb in Mp313E×Mp717 and <1 ppb in CML322) but increased by 5 and 14-fold in infested ears of resistant hybrid and inbred respectively. This suggested that the resistance to *A. flavus* colonization and AF contamination might have been compromised to some extent in seeds heavily damaged by CEW.

**Fig. 2.**
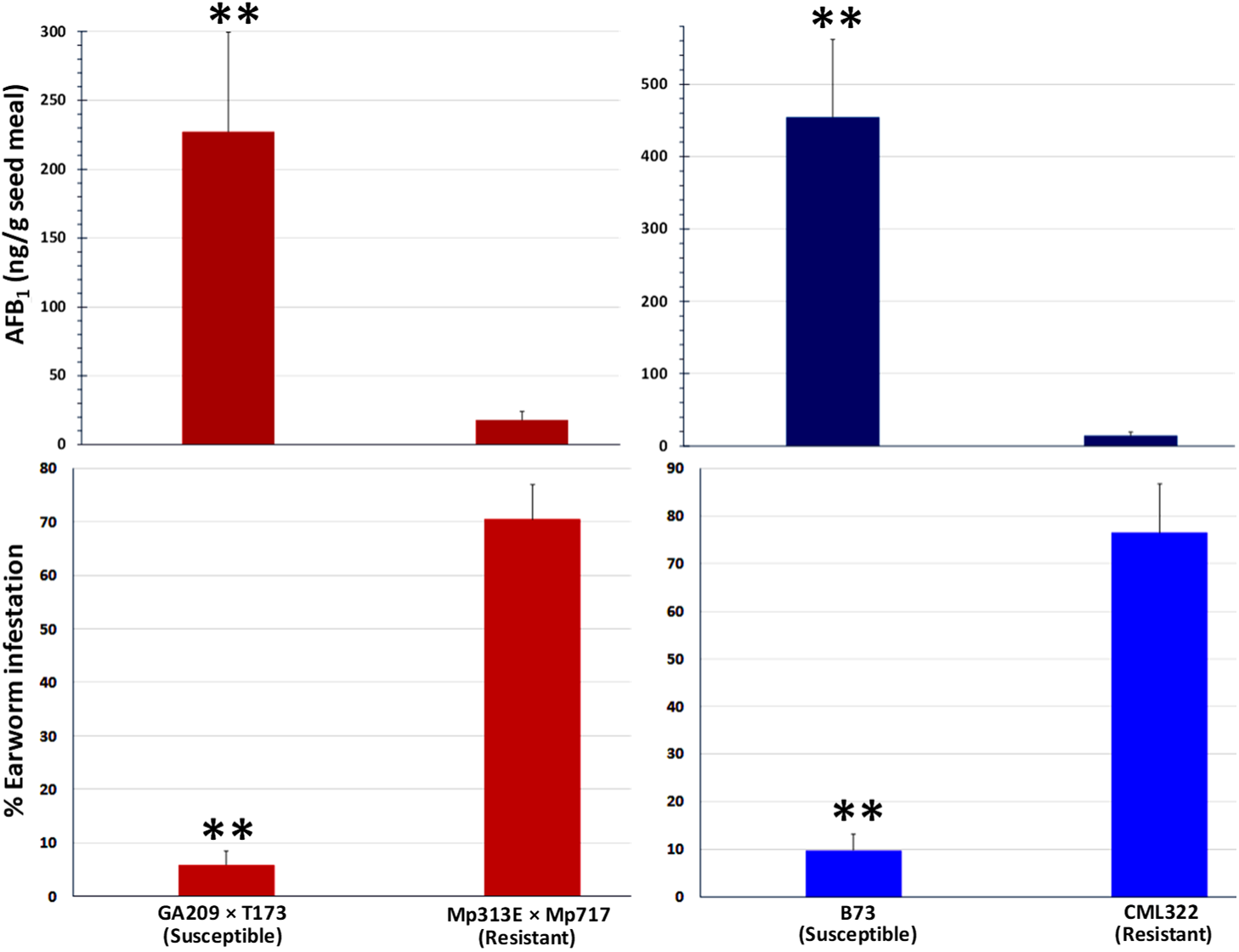
CEW damage is negatively correlated with seed AF content in maize lines. The infestation and AF data from control and infected ears is combined in each genotype. Significant differences (P value <0.05) between each data set were tested using an ANOVA (Supplemental Table 1) followed by Tukey’s multiple-comparisons post hoc test (Supplemental Table 2) in R (version 3.6.2). Average (+SE) infestation and AF values between *A. flavus* susceptible and resistant lines are highly significant (p<0.01).

### Kernel fumonisin content was enhanced in CEW-infested ears

*Fusarium verticillioides* is among the most common mycotoxigenic fungi colonizing field-grown maize. We observed symptoms of *F. verticillioides* colonization (e.g., star-burst pattern on seeds) in our samples. We isolated the fungus from seeds with visual symptoms using *Fusarium*-selective Malachite Green Agar 2.5 medium (Alborch et al. 2009) and confirmed by genomic PCR using *F*. *verticillioides*-specific primers (Baird et al. 2008). FUM content was analyzed in the same seed samples used for AF determination (**Fig. 3 A**) and compared between uninfested and CEW-infested samples (**Fig. 3B**).

**Fig. 3.**
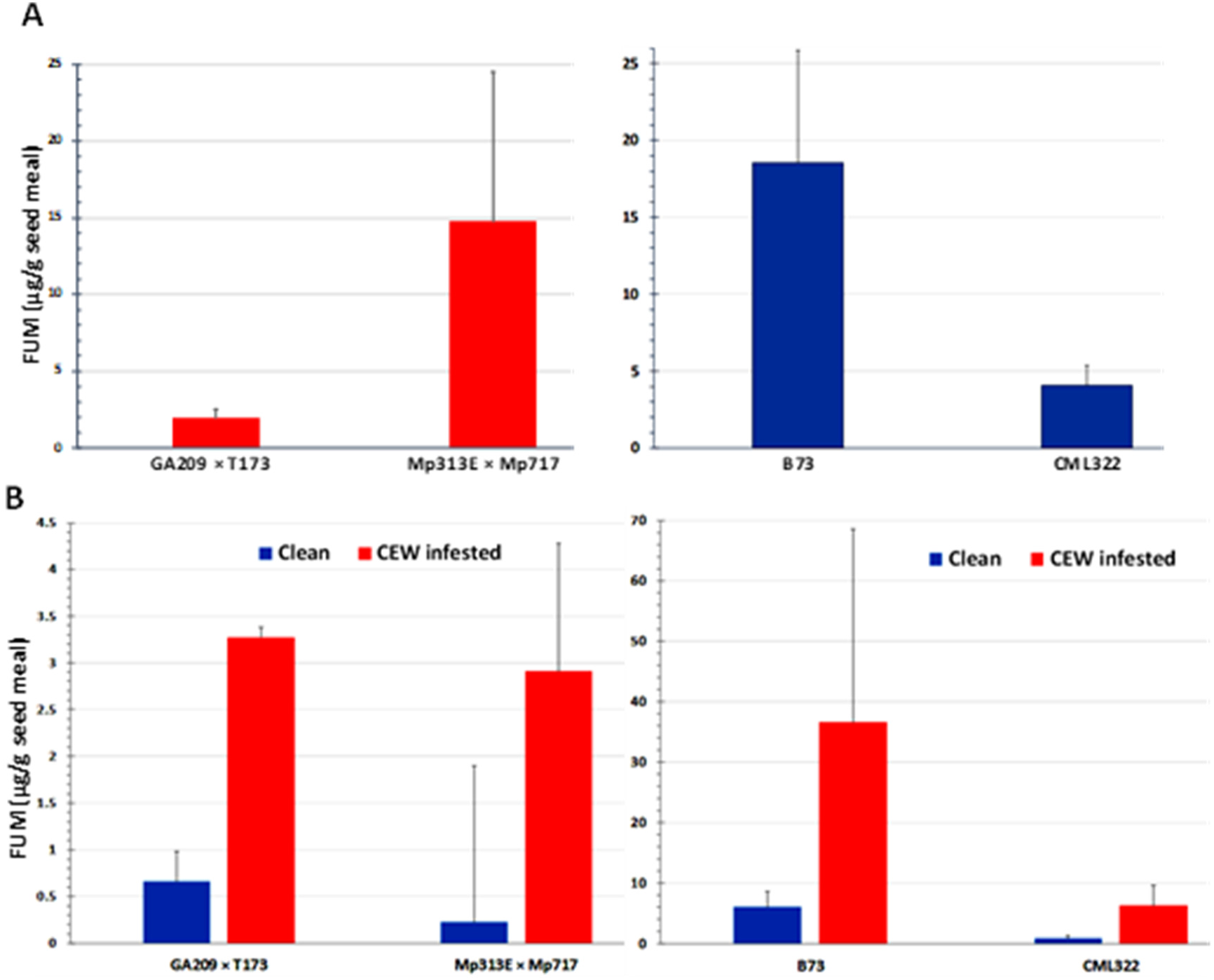
FUM contamination by native *Fusarium* strains. (**A**) Seed fumonisin content in the four maize lines. (**B**) Seed FUM content parsed by uninfested (clean) versus CEW infested ears in each genotype. The values are averages + SE in each genotype and were not significantly different at 95% confidence level.

Both maize hybrids used in this study have been previously shown to be resistant to FUM accumulation. In particular, Mp313ExMp717 (*A. flavus* resistant hybrid) was shown to be more robustly resistant than GA209xT173 across studies (Williams 2006; Henry et al. 2009; Williams and Windham 2009). In the current study, however, the Mp313ExMp717 accumulated >7-fold FUM in its seeds than GA209xT173 (**Fig. 3A**). Although CML322 accumulated a considerable amount of FUM, it was >4-fold less than that in B73, which is known to be among the most susceptible inbreds to Fusarium ear rot and FUM accumulation (Morales et al. 2019). However, when the data was parsed based on CEW infestation (only in sets where both clean and infested ears were available), infested ears showed >5-fold more FUM than uninfested ears (**Fig. 3B**). The differences were not significant probably due to the high variability in the colonization by native strains (the lowest p-value was 0.052 for CML322; also see **Fig. S2**). These data indicated that CEW may vector *Fusarium* spp. that produce FUM during its infestation.

### Differential toxicity of AF versus FB1 to CEW

The preferential infestation of *A. flavus* resistant lines by CEW and a negative correlation between AF and infestation rate, taken together with greater FUM levels in infested ears, suggested that AF may be more toxic to *H. zea* than FUM. We tested this hypothesis by feeding experiments where CEW neonates were reared on artificial diet containing graded levels of AF or FB1. Results shown in **Fig. 4** and **5** clearly demonstrate that the pest is more susceptible to AF than to FB1. As reported previously (Zeng et al. 2006), AF retarded CEW larval growth even at the lowest concentration tested, although the effect was not significant (**Fig. 5**) and was toxic above 200 ppb (**Fig. 4**). On the other hand, FB1 was non-toxic to CEW even at the highest concentration tested. In fact, at lower concentrations (below 30 ppm) the toxin seems to marginally enhance the growth of the larvae (the effect was consistent although there was variability among the bioassays). These results further support the proposal that the enhanced infestation of *A. flavus* resistant maize lines by *H. zea* may be due to very low levels of AF that are not inhibitory to larval growth.

**Fig. 4.**
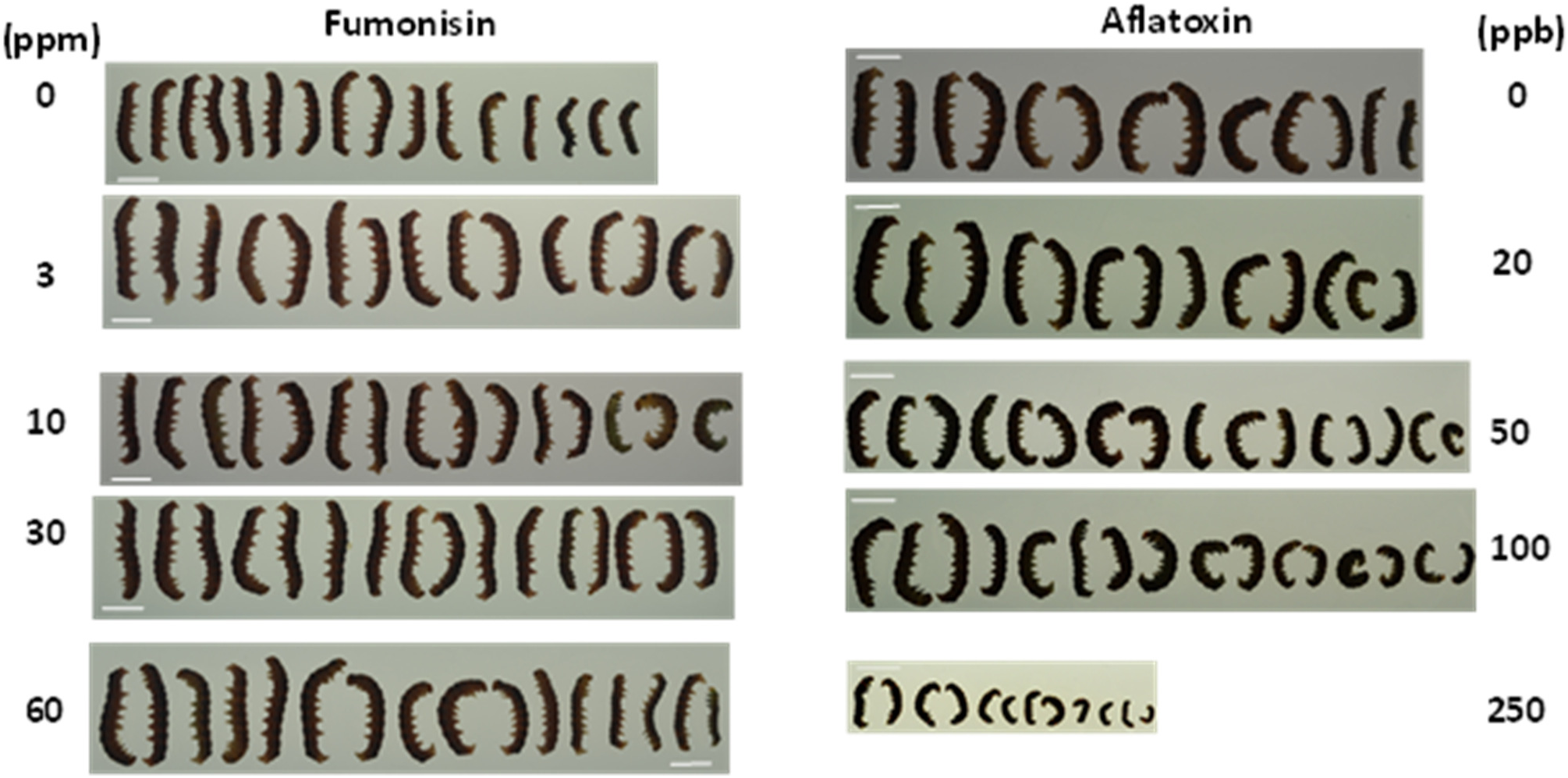
Effect of aflatoxin B_1_ and fumonisin B_1_ on the growth and mortality of *H. zea* larvae. Graded doses of AF or FB_1_ was tested on CEW growth and mortality by incorporating them into an artificial insect diet. Larvae were grown in a 128 well bioassay plate for 10 d. Each well had 1 g of feed and a single neonate at the start of the assay. A representative assay from 4 replicates is shown. In an additional assay, 100 ppm of FB_1_ and 300 ppb of AF were tested. Results were not different, except for a greater larval mortality at 300 ppb of AF (data not shown). Scale Bar = 1 cm.

**Fig. 5.**
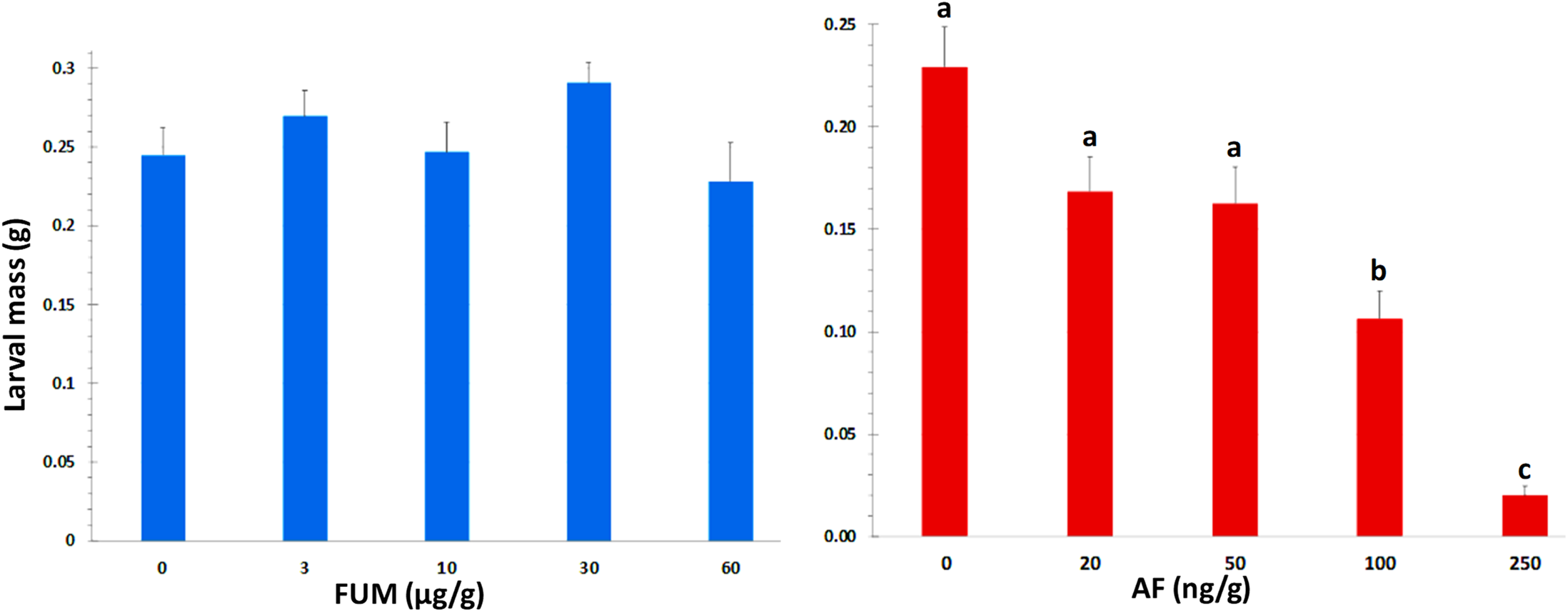
AF and FB_1_ effects on CEW larval mass. At the end of the bioassay, larvae were removed from the well killed by chloroform vapors and weighed. Values are averages + SE of ≥16 larvae/treatment except at 250 ppb of AF, where mortality was 30% or greater (dead and dried were seen stuck to the bottom of the well). The values marked with the same letter are not statistically significant. FB_1_ had no significant effect on larval growth at concentrations tested.

### Delayed flowering in *A. flavus* resistant maize lines

The tassel and ear development were delayed in CML322 by 3 weeks relative to B73 and by 4-5 weeks in the resistant hybrid, Mp313E×Mp717 compared to GA209×T173, although all four lines were planted together. CML322 is a tropical inbred and shows delayed flowering under long days, i.e., ≥13 h photoperiod (Hung et al. 2012). The parents of the resistant hybrid (Mp313Ex Mp717) are also derived from the tropical maize race Tuxpeño (Scott and Zummo 1990; Williams and Windham 2006) and known to show late-flowering phenotype. This is true for most maize lines that are resistant to *A. flavus* and attempts to segregate the two traits have been of limited success (Henry 2013). The availability of green silks may be an important factor for the increased *H. zea* infestation of these late flowering genotypes. However, in an adjacent plot where B73 was planted two weeks later (unrelated to the current study), silk emergence coincided with that of CML322 plants used in the present study. Nonetheless, B73 ears had highly elevated levels of seed AF (400 ppb in controls and 800 ppb in inoculated plants) and low levels of CEW infestation in this plot as well, suggesting that high seed AF levels may act as a deterrent for CEW infestation because of its toxicity.

## DISCUSSION

There are few studies where CEW infestation patterns have been compared in maize genotypes with varying resistance to *A. flavus* or AF accumulation. Nie et al (2011) compared spatial patterns of natural infestation of four ear-feeding insects (CEW, fall armyworm, maize weevil and brown stinkbug) with AF distribution due to colonization of a single commercial maize hybrid by native *A. flavus* strains. In the first year of the study, CEW infestation was very extensive (95% of ears) and in the second year, although less intense, it was as high as 41%. However, AF contamination was very low in both years (>80% of ears had ≤30 ppb and only ≤4% ears had ≤100 ppb). Although the predominantly low AF content makes it difficult to quantify the relationship, it is strongly indicative of a negative association between CEW damage and seed AF content. The maize genotypes in our study have proven resistance or susceptibility to *A. flavus*. Further, high AF contamination (≤100 ppb) in uninoculated as well as inoculated plots of only susceptible lines allowed to make robust comparisons.

The premise for this study is an unprecedented or unreported observation, in that two unrelated maize lines (Tuxpeño germplasm versus CML) with proven resistance to *A. flavus* were heavily infested by CEW. Conversely, the two *A. flavus* susceptible lines (stiff-stalk inbred B73 and non-stiff stalk hybrid GA209 x T173) were spared from heavy CEW damage. Although late flowering maize is known to be susceptible to CEW infestation by providing green silks, availability of silks alone could not fully explain our observations. Late flowering is more often a problem in the northeastern US where it coincides with CEW migration from southern states. Furthermore, late planted B73 in an adjacent plot had delayed silk emergence but showed no CEW infestation. The other and more likely explanation is that the susceptible lines had very high levels of AF that were toxic to CEW. Even mock-inoculated controls had 100 ng of AF per gram of seed meal prepared from entire ears with both moldy and non-moldy seeds. This argument is supported by previous studies on AF toxicity to CEW in feeding experiments (Zeng et al. 2006) as well as our current work (**Fig. 4** and **5**). Zeng et al (2006) showed that AF at 200 ppb strongly inhibited the growth and development of first instar larvae, leading to >50% larval death after 9 d and 100% death after 15 d of feeding. Even lower concentrations (1-20 ppb; FDA-regulated levels) affected larval development, delayed pupation rate and led to >40% mortality when the exposure was longer than 7 d (Zeng et al. 2006). Although concentrations below 20 ppb were not tested in our study, we observed a steady decline in larval mass as AF concentration increased with ≥30% mortality at or above 250 ppb during 10-15 d exposure (**Fig. 5**). We did not continue our observations beyond the larval stage to assess the longer term developmental effects (e.g., pupation or emergence of adults). An apparent exception to the correlation between low AF and high CEW infestation was a significant decrease in CEW infestation observed in TOX4-inoculated ears compared to uninoculated ears in the *A. flavus* resistant inbred CML322, although average AF levels did not exceed 30 ppb. Given the highly variable distribution of AF in individual kernels of a maize ear (e.g., Lee et al. 1980), it is possible that AF content particularly in damaged kernels (close to the silk canal, the site of inoculation as well as CEW infestation) was much greater than the average for the entire ear and high enough to be toxic to CEW survival. Furthermore, CEW may be sensitive also to other anti-insectan compounds made by *A. flavus* (Cary et al. 2018) that could act additively or synergistically with AF (e.g., Kojic acid; Dowd 1988). Future experiments would involve late-maturing lines with *A. flavus* susceptibility and early maturing lines with *A. flavus* resistance to clarify and quantify the effects of flowering time and AF content on CEW infestation.

It is not surprising that AF is toxic to insects, not merely to mammals. *A. flavus* is predominantly a soil-living saprophyte, feeding on decaying organic matter, including dead insects. It is also an opportunistic pathogen and can colonize a wide variety of insects, e.g., moths, silkworms, bees, grasshoppers, houseflies and mealy bugs among others (St. Leger et al. 2000; Gupta and Gopal 2002 and references therein). At the same time, *A. flavus* is known to survive ingestion by mycophagous insects. Among three *Aspergillus* species tested, *A. flavus* conidia phagocytized by insect hemocytes were still able to germinate (St. Leger et al. 2000). *A. flavus* may also proliferate in the hindgut of CEW (Abel et al., 2002). In spite of being a polyphagous pest with a remarkable capacity to metabolize a wide array of plant compounds, CEW has limited tolerance to AF and poor ability to metabolize the mycotoxin (Dowd 1988; Zeng et al. 2006). The fungus is known to make several anti-insectan compounds, beside AF (TePaske et al. 1992; Cary et al. 2018). Other insect pests that are more tolerant may vector *A. flavus* (Zeng et al. 2006; Opoku et al. 2019). Spatial correlation analysis of natural infestation by different pests and seed AF content in field-grown maize plants indicated that AF content was correlated to the frequency of weevils and stink bug-affected kernels, but not with CEW damage (Ni et al 2011).

Our work also showed that FUM is not toxic to *H. zea* (**Fig. 4**). This may have allowed CEW to vector *F. verticillioides* and other FUM-contaminating fungi, as indicated by an increased seed FUM content in infested ears (**Fig. 3**). CEW damage is also frequently associated with the colonization by another mycotoxigenic fungus, *Stenocarpella maydis*, which causes diplodia ear rot (Munkvold and White 2016). In animal model systems, FB1 at 25-50 µM (i.e., 18-36 ppm) inhibits ceramide synthases and leads to the accumulation of toxigenic/carcinogenic sphinganine and related compounds (Riley et al., 2001; Riley and Merrill 2019). Conversely, FB1 was not toxic to yellow mealworm larvae even at 450 ppm when included in the diet or when injected into larva (Abado-Becognee et al. 1998). Recently, the brown marmorated stink bug (*Halyomorpha halys*) was shown to enhance *F. verticillioides* infection and FUM contamination in field corn (Opoku et al. 2019). Among other secondary metabolites produced by *F. verticillioides*, fusaric acid is only a weak antisectan compound (Dowd 1988). The lack of secondary metabolites with potent insecticidal properties in the biosynthetic repertoire of *F. verticillioides* could be one of the reasons for its frequently observed transmission via insect infestation (e.g., Smeltzer 1959; Dowd 2000; Mesterházy et al. 2012; Madege et al. 2018)

The association between CEW-infestation and high FUM content can also be explained by host reaction to fungal infection potentially triggering enhanced insect damage. Mycotoxin-producing *Fusarium* spp. trigger volatile production by maize leaves that attract cereal leaf beetles (Piesik et al. 2011). Other examples where insect species benefit from the presence of mycotoxigenic fungi are also reported (Schulthess et al. 2002). Alternatively, insect-fungus interactions can enhance production of secondary metabolites by plant host tissues (Döll et al. 2013; Drakulic et al. 2015, 2016).

Although this study was pursued to explain a serendipitous observation made during two unrelated field studies, it has important implications in mycotoxin control. AF and FUM are ubiquitous and unpredictable contaminants of commodities, particularly maize. Our study clarifies a component of this unpredictability. The late flowering trait of *A. flavus* resistant lines (owing to their tropical origin) is known to delay harvest, potentially leading to frost damage and/or high grain moisture. Our current work shows that delayed flowering coupled with low AF accumulation can exacerbate CEW infestation, which in turn can lead to contamination by other mycotoxins, such as fumonisins (Munkvold and White 2016).

In contrast to a mutual antagonism reported previously between *A. flavus* and *F. verticillioides* (Zummo and Scott 1992; also see **Fig. S3**), we observed high levels of AF and FUM co-contaminating our samples. B73, in particular with its high susceptibility to both mycotoxigenic fungi, had very high levels of both AF and FUM in many of its seed samples. Although CEW damage was very low in this inbred (**Fig. 1B and 2**), FUM levels were exacerbated in infested ears (**Fig. 3B**). There is some evidence for an additive or even synergistic effect on carcinogenicity from co-exposure to AF and FUM (World Health Organization 2018). Based on biomarker studies and food analyses, the co-occurrence of these two mycotoxins has been widely documented in developing countries (Shirima et al. 2013; Biomin Mycotoxin Survey 2019). It is important to examine the underlying factors as well as effects of mycotoxin co-contamination both by researchers and regulatory agencies to mitigate its impact on food safety (Lopez Garcia 1998).

## MATERIALS AND METHODS

### Field planting of maize and application of *A. flavus* toxigenic strains

The four maize genotypes used in the study are non-transgenic and non-commercial lines. The two hybrids, GA209×T173 (susceptible to AF accumulation) and Mp313E×Mp717 (resistant to AF accumulation), were developed at the USDA-ARS Corn Host Plant Resistance Research Unit, Mississippi (Williams and Windham 2009). The hybrids, along with two popular inbreds B73 (susceptible to AF accumulation, (Campbell and White 1995) and CML322 (resistant to AF accumulation, (Betrán et al 2002)) were planted in 4-row plots at the LSU Agricultural Experimental Station in Baton Rouge (Louisiana) in the middle of April. To keep the insect pressure low, Besiege (a broad-spectrum foliar insecticide with fast knockdown and long-lasting residual effects; has chlorantraniliprole and λ-cyhalothrin as active ingredients) was sprayed at ∼V9 and R1 growth stages. Three days after the second insecticide application, plants were inoculated with *A. flavus* strains by silk canal injections (Zummo and Scott 1992), with conidial suspensions as described before (Chalivendra et al. 2018). The hybrids were inoculated with CA14, inbreds with Tox4. Plants were maintained with standard agronomic practices of fertilizer and herbicide applications and received irrigations during extended dry periods.

The inbred study was originally aimed at analyzing microbiome changes in a susceptible and a resistant line in response to *A. flavus* colonization. We used Tox4 in the study because it is an isolate from local maize fields (Chalivendra et al. 2018), produces high AF levels and serves as a good model strain to study microbiome changes. The experiment with hybrids was an extension of recent studies on biofilm-like structure formation by *A. flavus* during maize seed colonization (Dolezal et al. 2013; Shu et al. 2014; Windham et al. 2018). The objective of our study was to localize the expression of *A. flavus* Medusa A gene by *in situ* hybridization in maize seeds in relation to the spatial distribution of the biofilm-like structure. *A. flavus* strain CA14 was used in the study, since it has whole genome sequence information and needed mutant resources (Chang et al. 2019). CA14 was obtained from the USDA Agricultural Research Service Culture Collection, Northern Regional Research Laboratory, Peoria, IL, USA.

### HPLC analysis of aflatoxin B_1_

One ear per plant from each genotype and treatment was harvested, resulting in 70-80 ears in inoculated plants and double the number from uninoculated plants. Ears in each lot were separated by the presence or absence of CEW infestation to monitor the effect of insect damage on mycotoxin levels. Only ears with visible internal damage (i.e., nibbled seed and cut silks, larval feeding tracks with frass; sometimes with dead or live CEW larvae) were considered as infested. No distinct spatial or other pattern of infestation was observed in our plots (as was also reported by Ni et al. 2011), except that a majority of resistant inbred or hybrid plants were infested, while only a few ears from susceptible lines showed damage by the earworm. At least three ears were used per replicate and each category had 3-5 replicates. Given the low frequency of CEW-damaged ears in B73 and GA209×T173, all ears in each category were used for AF analysis to have robust AF data. When the seed meal exceeded more than 100 g (in uninoculated controls), we took more than one sample to minimize sampling error. AF from seed meal was extracted and measured as before (Chalivendra et al. 2018) with modified HPLC conditions. The equipment included Waters e2695 HPLC (Waters Corp., Milford, MA, United States) fitted with a Nova-Pak C18 column, a photochemical reactor (Aura Industries Inc., New York, United States) and a Waters 2475 FLR Detector (Waters Corp.). The signal was detected by excitation at 365 nm and emission at 440 nm. Aqueous methanol (37.5%) was used as the mobile phase.

### LC-MS analysis of fumonisins

Maize kernel samples were analyzed for FB1, FB2 and FB3 by liquid chromatography–mass spectrometry (LC-MS) using an adaptation of a previously published method for mycotoxin analysis (Plattner 1999). Briefly, maize samples were ground with a laboratory mill. Portions (5 g) of the seed meal were extracted with 25 mL 1:1 acetonitrile/water for 2 h on a Model G2 Gyrotory Shaker (New Brunswick Scientific, Edison, NJ, USA). Extracts were filtered with a Whatman 125 mm 2V paper filter (GE Healthcare Bio-Sciences, Pittsburgh, PA, USA). A total of 10 µL of extract was applied to a Kinetex (Phenomenex, Torrance, CA, USA) C18 column (50 mm length, 2.1 mm diameter). Chromatography was conducted utilizing a Thermo Dionex Ultimate 3000 (Thermo Fisher, Waltham, MA, USA) ultrahigh-performance liquid chromatography (UPLC) system consisting of an autosampler coupled to a binary gradient pump. Elution of analyte was achieved with a 0.6 mL min−1 gradient flow of methanol and water (0.3% acetic acid was added to the mobile phase). The solvent program used a 35–95% gradient over 5 min. Flow was directed to a Q Exactive (Thermo Fisher, Waltham, MA, USA) hybrid quadrupole-Orbitrap mass spectrometer equipped with an electrospray ionization source. The mass spectrometer was operated in full-scan mode over a range of 300 to 1200 m/z. Operation of the LC-MS and quantification of the eluting fumonisins were performed utilizing Thermo Xcalibur software. Quantification of fumonisins was based upon intensity of protonated ions for FB1 (m/z 722.3), FB2 (m/z 706.3) and FB3 (m/z 706.3) compared to calibration standards of the toxins. The limit of quantification for the analytical method was determined to be 0.1 µg per g for FB1, FB2 and FB3.

### Toxicity bioassays

The toxicity of FUM to CEW larvae was tested in a pre-mixed meridic diet (WARD’S Stonefly Heliothis diet, Rochester, NY) containing 0. 3, 10, 30 60 or 100 μg/g FB1 (Cayman Chemical, MI) or 20, 50, 100, 250 or 500 ng/g of AFB_1_ (Sigma Chemicals). The diet was prepared as per manufacturer’s instructions. The FB1 stock, made in water, was diluted to the above rates before the dry diet was added and mixed thoroughly. AF was dissolved in methanol at a stock concentration of 2 mg/mL and diluted appropriately to provide the aforementioned concentrations. The highest concentration of methanol used (0.08% by w/w) was incorporated into the control diet. The assay was done in a 128 well bioassay plate (C-D International Inc., Pitman, NJ). A single CEW neonate from a laboratory CEW colony obtained from Benzon Research Inc. (Carlisle, PA) was added to each well with 1 g diet using a camel hair brush (Kaur et al. 2019). At least 20 larvae were tested per treatment and the assay was repeated four times.

### Statistical analysis of data

Insect damage and aflatoxin levels were compared by ANOVA and post-hoc analysis by Tukey’s Honestly Significant Difference (HSD) test using R program (version 3.6.2) in RStudio. Student’s t-test was used for comparison of specific pairs of data sets.

### Safety

Aflatoxin B_1_ and fumonisin B_1_, being highly toxic mycotoxins, were handled with care using a biohood, surgical gloves and nose as well as mouth masks. All residues and containers were decontaminated using bleach and by autoclaving.

## ACKNOWLEDGMENTS

SC thanks the National Corn Growers Association for the funding support through their AMCOE program. Excellent field and laboratory assistance by Mr. Anthony Nguyen is gratefully acknowledged. Dr. Z-Y Chen is thanked for his help in aflatoxin measurement.

## Supplemental data

**Figure S1.**
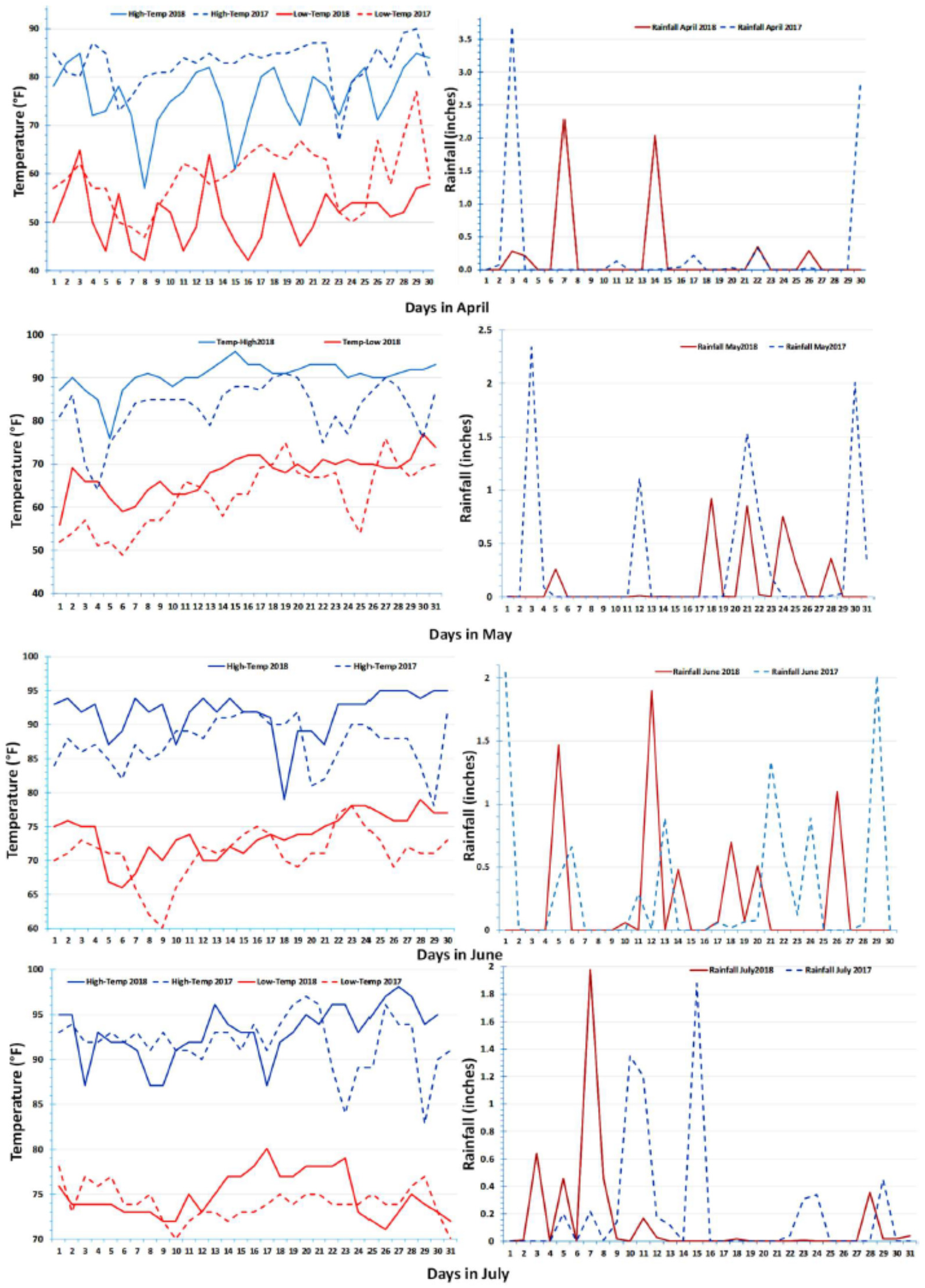
Weather data for the months of April-July in 2018 (solid lines) and 2017 (dashed lines). Daily high (blue lines) and low (red lines) temperatures are shown in the left panel. Rain fall is shown in the right panel.

**Fig. S2.**
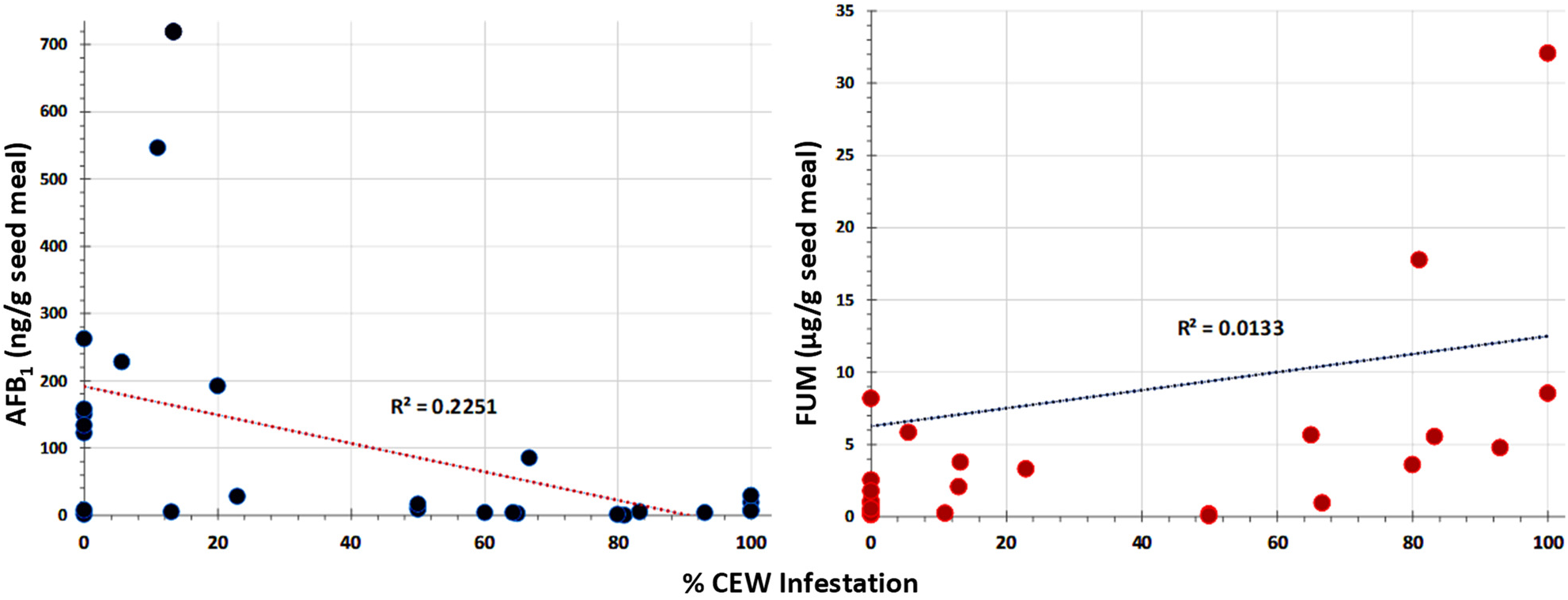
Correlation between CEW infestation of ears and seed AF or FUM levels in maize. Combined data from inbred and hybrid maize lines is plotted. CEW showed a negative relationship with AF and a positive trend with FUM. The greater correlation observed with AF (Pearson correlation coefficient, R = −0.47) was likely because of manual inoculation with specific strains of *A. flavus* (dominant to native strains), whereas more random infestation by native *Fusarium* strains may have led to poor correlation (R = 0.115).

**Fig. S3.**
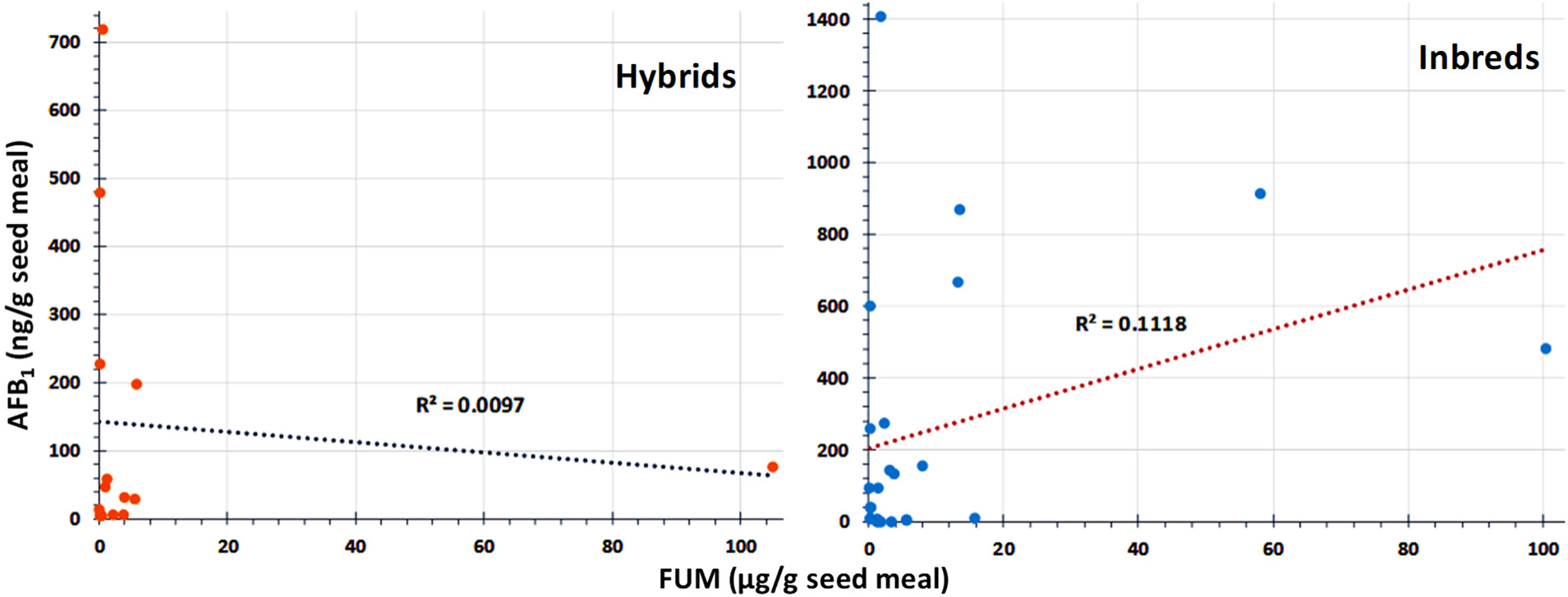
Correlation of Seed FUM and AF contents in hybrids and inbreds. Contents of the two mycotoxins from the same seed sample are poorly correlated in both sets as indicated by Pearson correlation coefficient values (r = −0.0983 for hybrids and 0.3344 for inbreds). This lack of correlation indicated that there was no mutual effect in the production of the two mycotoxins by the fungi infecting seeds from same ears.

**Table S1.**
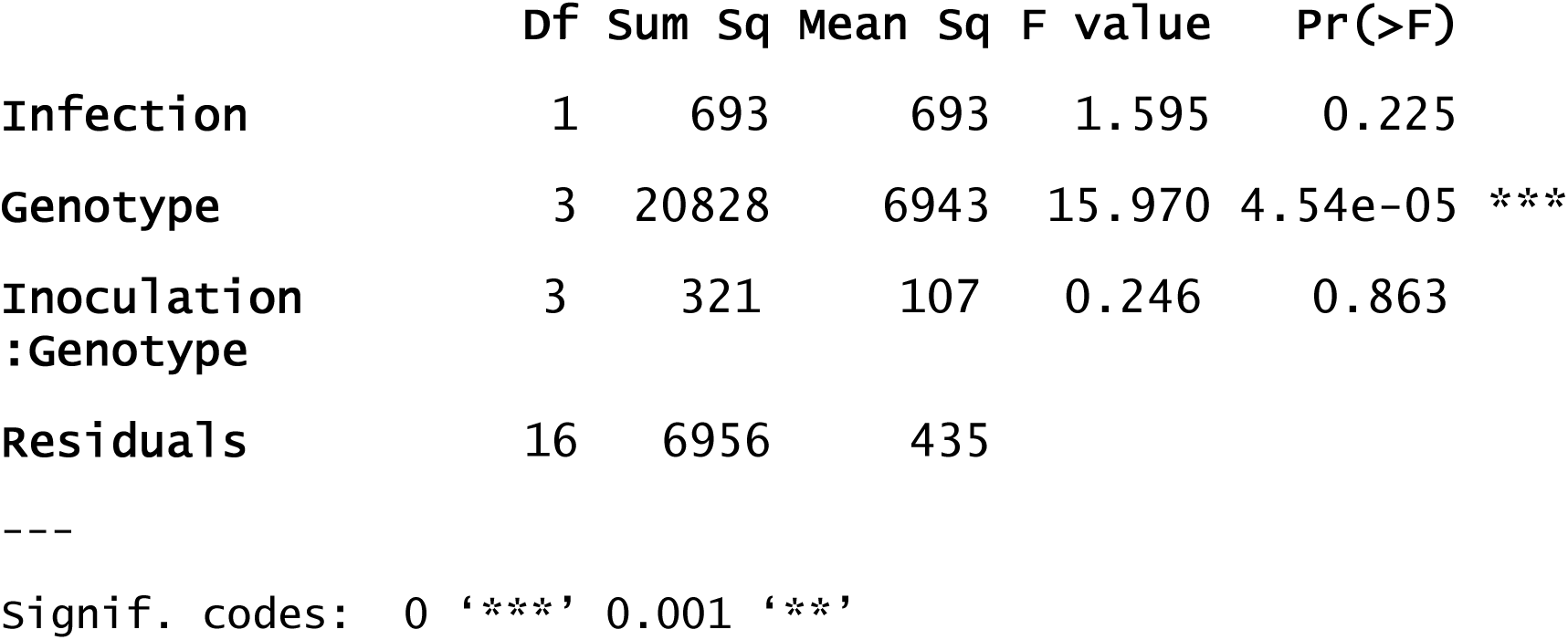
Analysis of variance for CEW infestation in maize inbreds and hybrids with differential resistance to aspergillus ear rot.

|  | **Df** | **Sum Sq** | **Mean Sq** | **F value** | **Pr(>F)** |
| --- | --- | --- | --- | --- | --- |
| **Infection** | 1 | 693 | 693 | 1.595 | 0.225 |
| **Genotype** | 3 | 20828 | 6943 | 15.970 | 4.54e-05 *** |
| **Inoculation**<br>**:Genotype** | 3 | 321 | 107 | 0.246 | 0.863 |
| **Residuals** | 16 | 6956 | 435 |  |  |

Signif. codes: 0 '\*\*\*' 0.001 '\*\*'

**Table S2.**
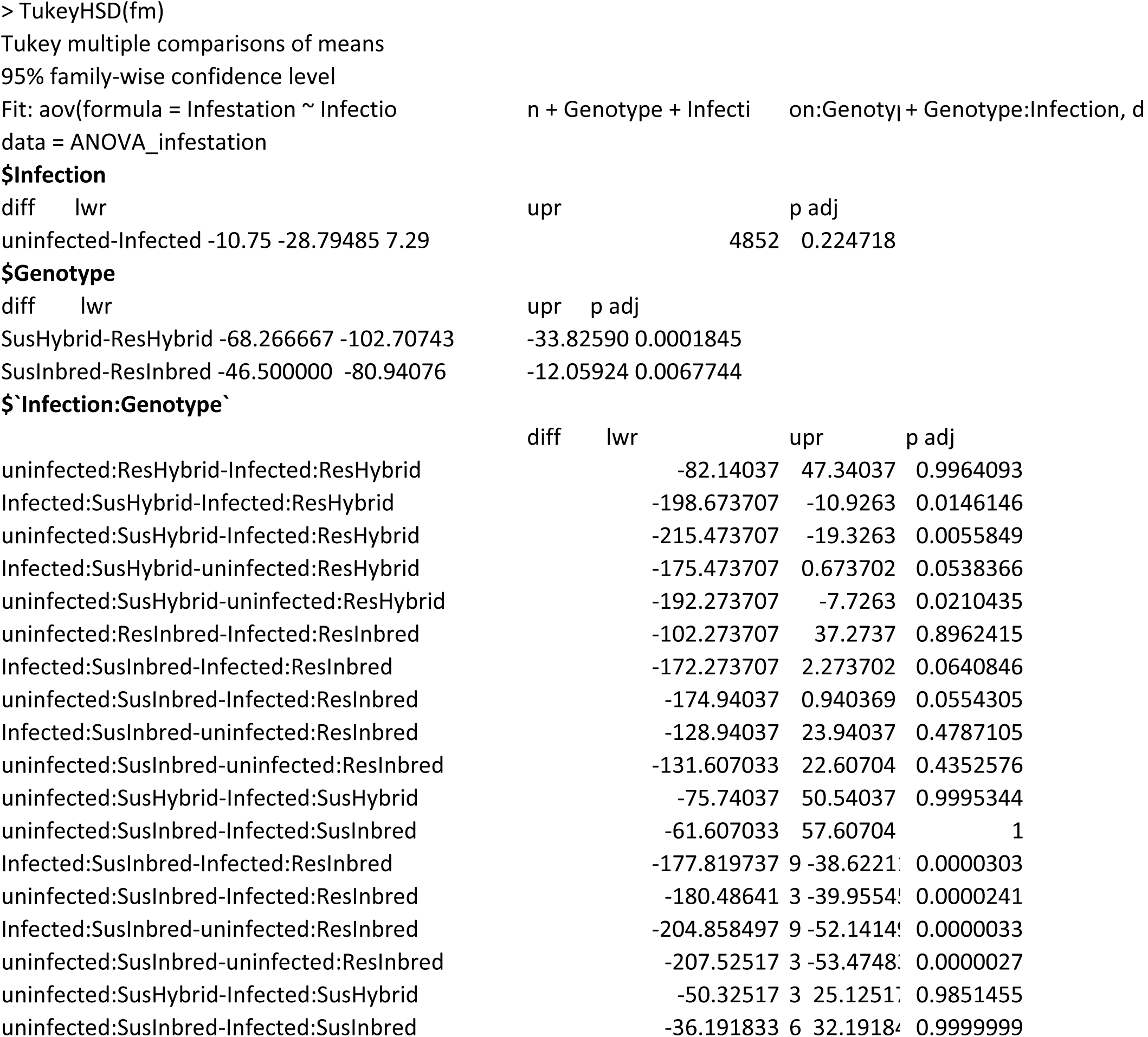
Tukey HSD for Infestation data.

**Table S3.** Analysis of variance for seed aflatoxin content in maize inbreds and hybrids with differential resistance to aspergillus ear rot.

|  | **Df** | **Sum Sq** | **Mean Sq** | **F value** | **Pr(>F)** |  |
| --- | --- | --- | --- | --- | --- | --- |
| **Genotype** | 3 | 3623512 | 1207837 | 34.373 | 3.99e-10 | *** |
| **Infection** | 1 | 726043 | 726043 | 20.662 | 7.41e-05 | *** |
| **Infestation** | 1 | 308549 | 308549 | 8.781 | 0.005705 | ** |
| **Genotype:Inoculation** | 3 | 765751 | 255250 | 7.264 | 0.000753 | *** |
| **Genotype:Infestation** | 3 | 423363 | 141121 | 4.016 | 0.015605 | * |
| **Inoculation:Infestation** | 1 | 1979 | 1979 | 0.056 | 0.813916 |  |
| **Genotype:Inoculation**<br>**:Infestation** | 3 | 125570 | 41857 | 1.191 | 0.328698 |  |
| **Residuals** | 32 | 1124456 | 35139 |  |  |  |

Signif. codes: 0 '\*\*\*' 0.001 '\*\*' 0.01 '\*' 0.05 '.'

**Table S4.**
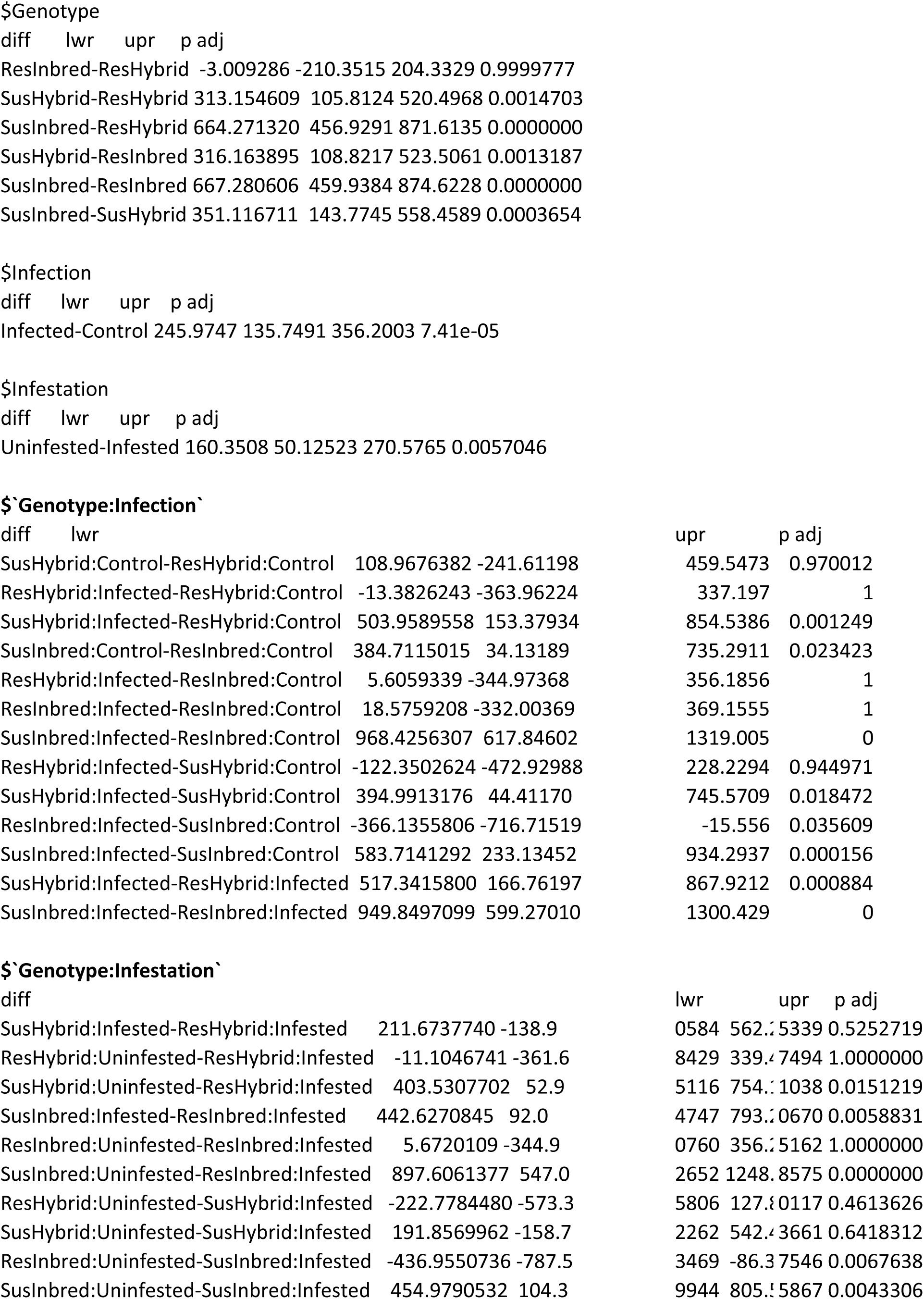
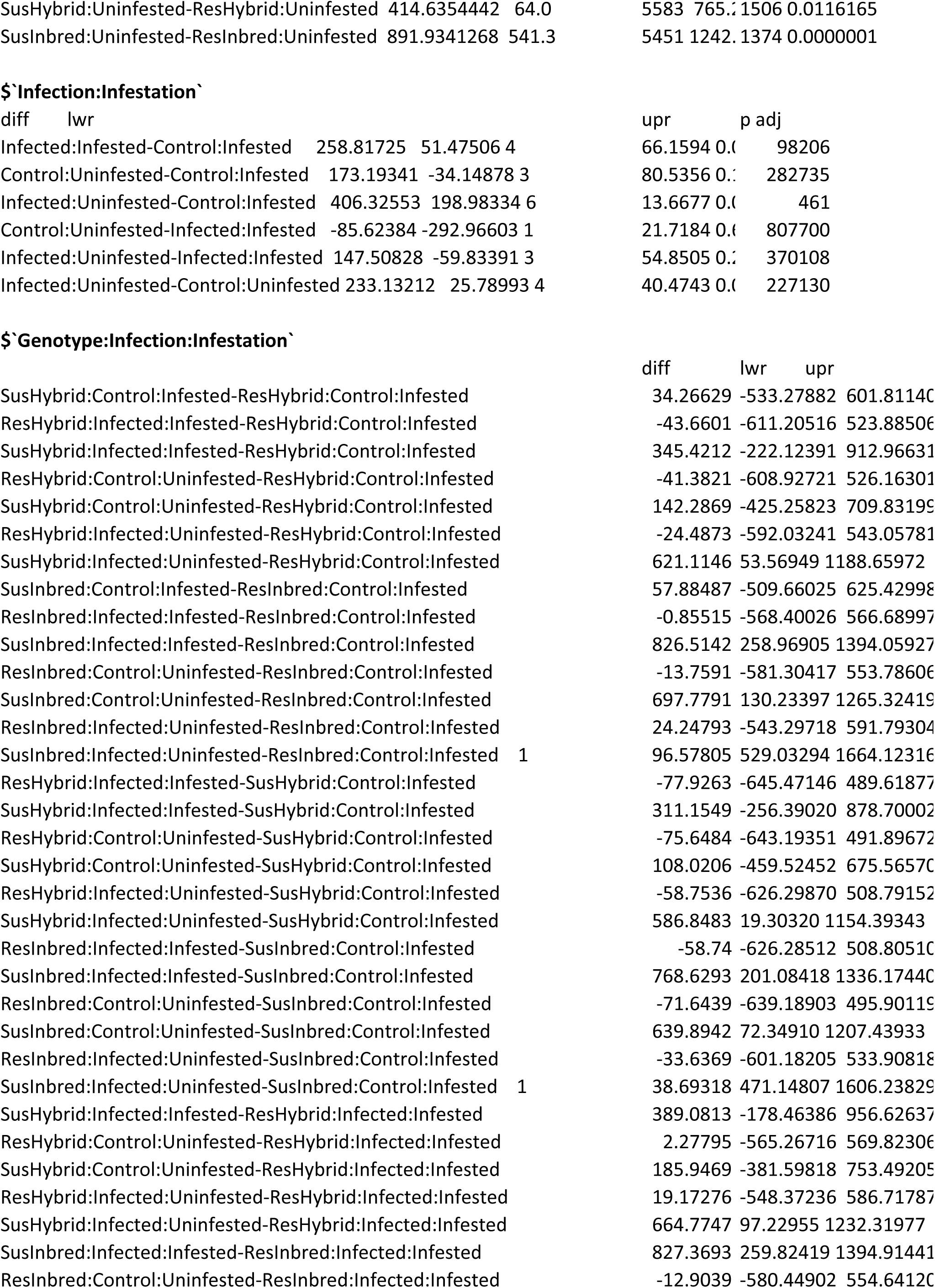
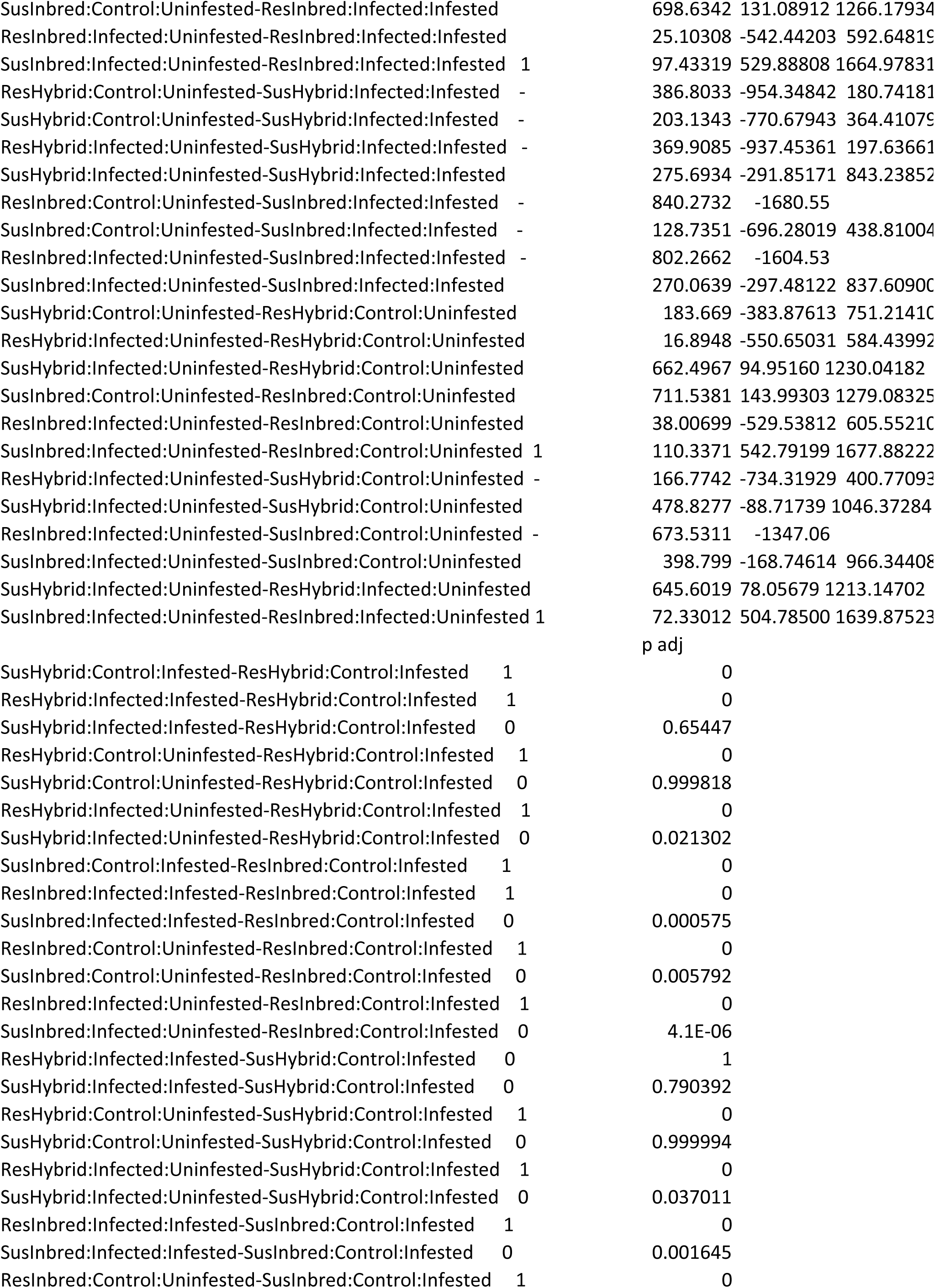
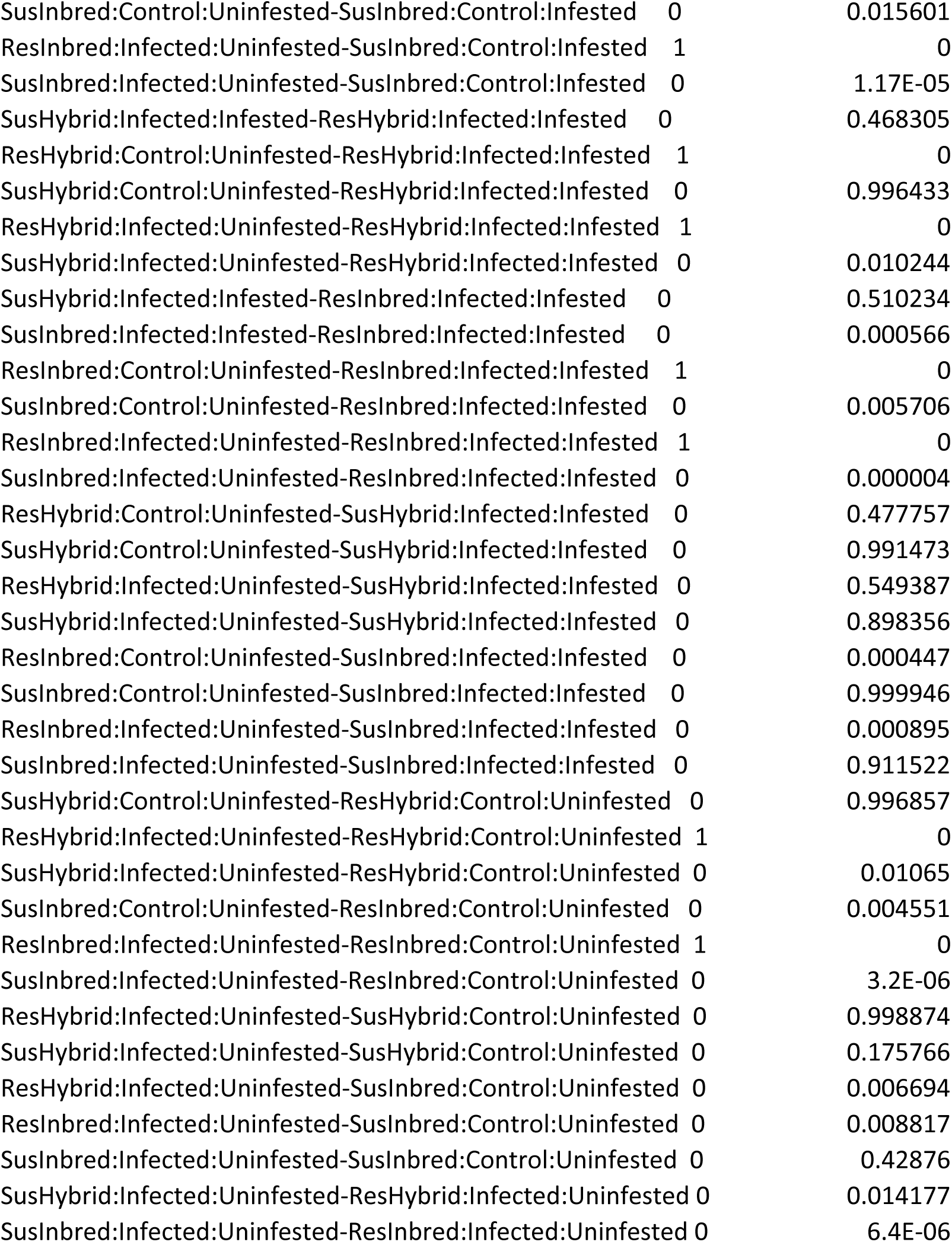
Tukey’s HSD for AF data.

